## Supplemental tables and figures for "Genetic Loci and Metabolic States Associated With Murine Epigenetic Aging"

**Supplementary Figures**

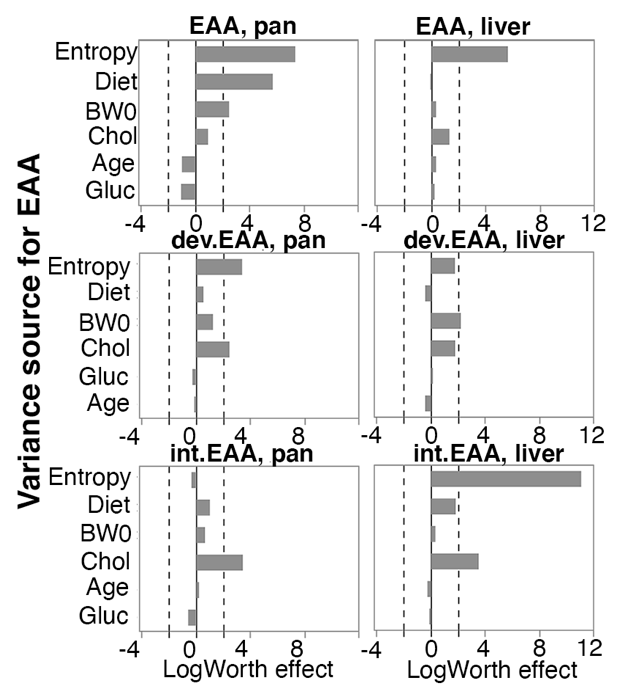

**Fig S1. Relative effects of different predictor variables on epigenetic age acceleration (EAA)**

Logworth scores of the predictors (–log_10_p) with dashed lines corresponding to p = 0.01. Positive logworth values indicate positive regression estimates, and negative values indicate negative regression estimates (for diet, positive means higher in high fat diet compared to control diet). BW0 is baseline weight; Chol is serum total cholesterol, Gluc is fasted glucose levels.

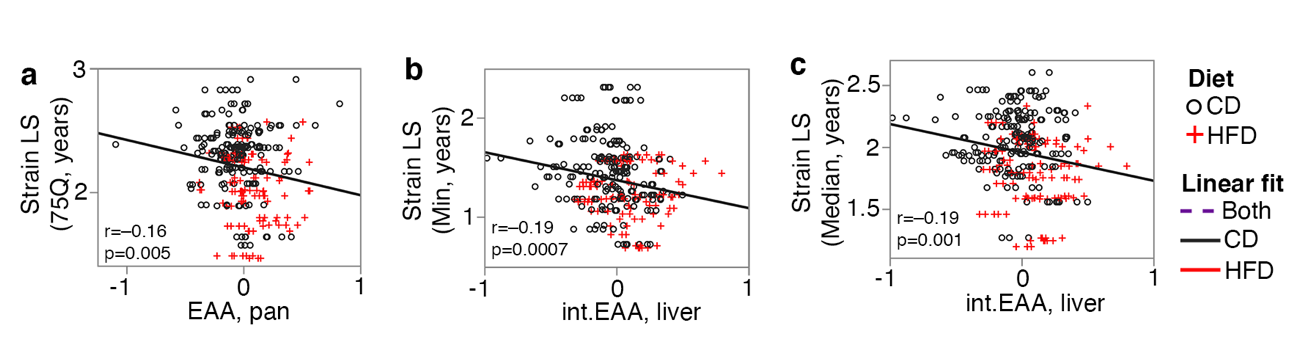

**Fig S2.** **BXD strains with shorter life expectancy have slightly more accelerated clocks.** This inverse correlation is depicted for the **(a)** 75^th^ quartile age at natural death, **(b)** the minimum lifespan, and **(c)** the median lifespan (analysis in 302 female samples with lifespan data). CD is control diet; HFD is high fat diet.

The negative correlations are modest with explained variance values, r^2^, of about ~3%.

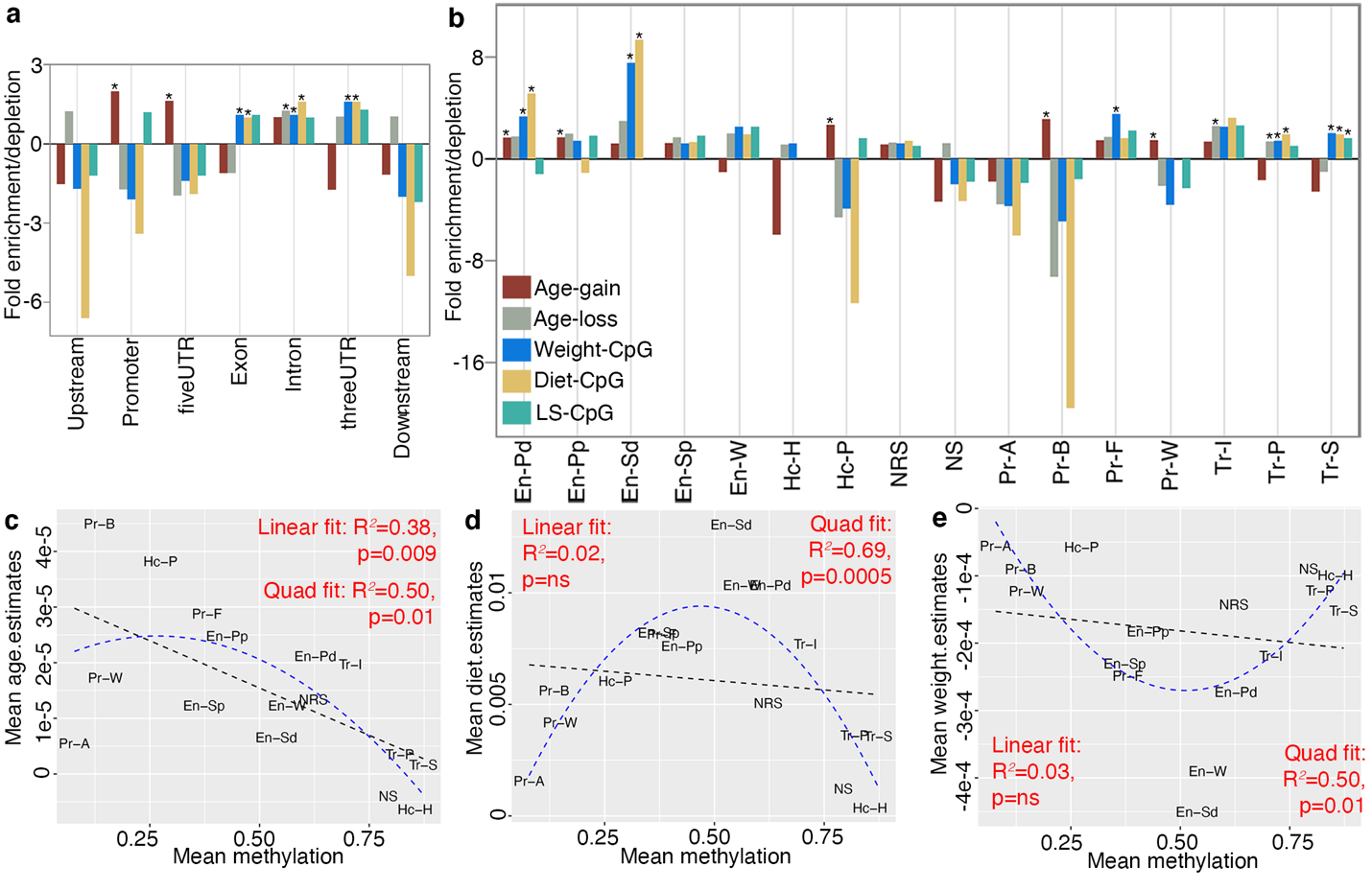

**Fig S3. Genomic and chromatin states of differentially methylated CpGs**

Enrichment is **(a)** genomic location, and **(b)** chromatin states among the differentially methylated CpGs (DMC) (expansions for the chromatin states are provided in **Data S8**). Asterisks denote hypergeometric enrichment p < 0.001. For the 15 chromatin states (and regions with no replicable signal, NRS), we compare the methylation levels, and mean regression estimates for the effects of **(c)** age, **(d)** diet, and **(e)** body weight.

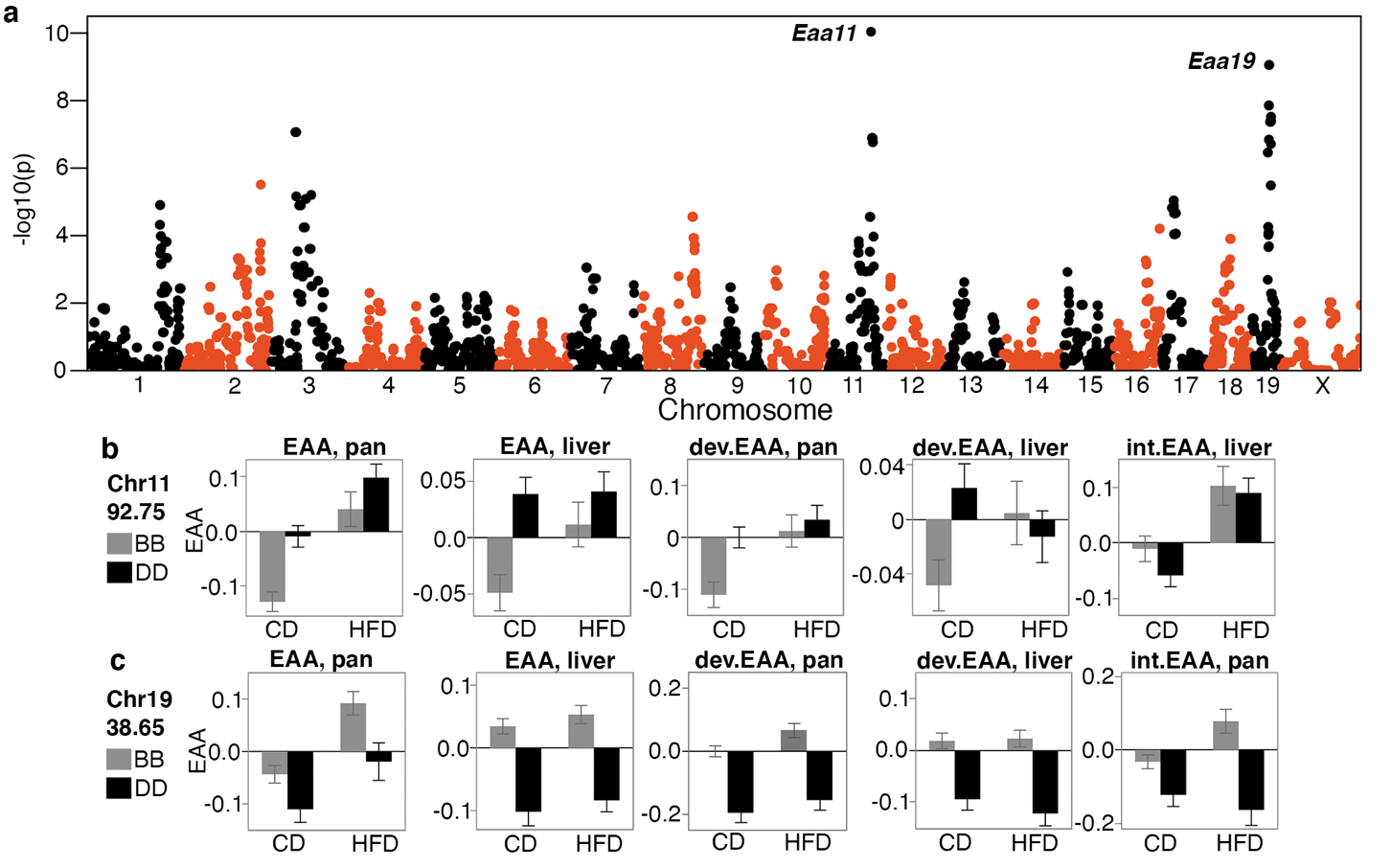

**Fig S4.** **Consensus QTL mapping for epigenetic age acceleration**

**(s)** The Manhattan plot displays the combined meta p-values for epigenetic age acceleration (EAA). These meta p-values are based on a simple p-value combination for the six EAA QTL traits, and is mainly to highlight regions with the highest consensus QTLs. The highest peaks are on chromosomes 11 (*Eaa11*), and 19 (*Eaa19*). **(b)** BXDs were segregated by genotype at a representative marker in *Eaa11* (variant at 92.750 Mb). In the control diet group (CD), mean EAA (± standard error) is higher for mice with the *DD* genotype. Only the EAA derived from the liver interventional clock (int.EAA) shows no difference between the genotypes. **(c)** BXDs were segregated by the genotype at a marker in *Eaa19* (38.650 Mb). Mean EAA is higher in the *BB* genotype, and this genotype effect is seen for all the clocks on both diets. Bars are standard error.

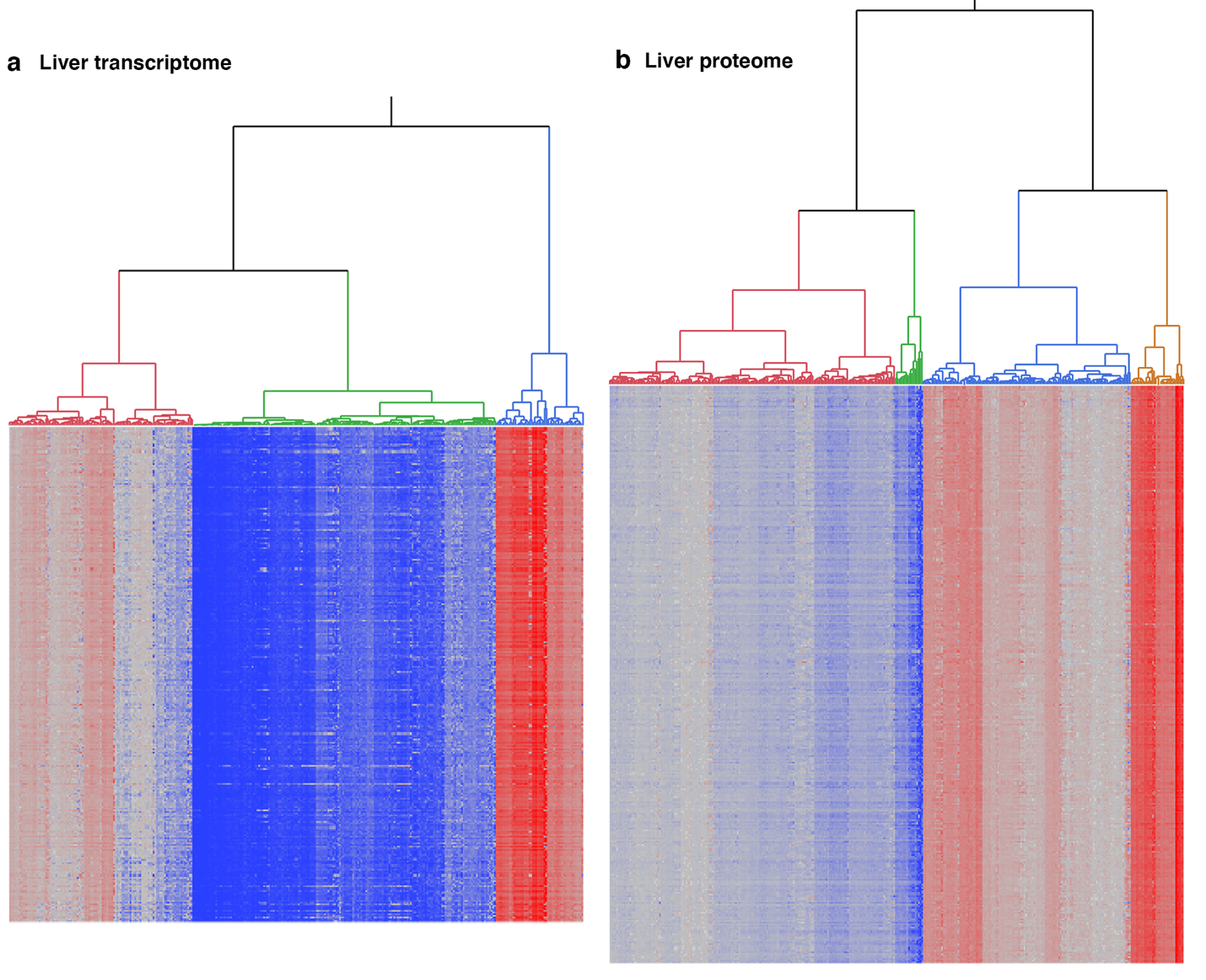

**Fig S5. Hierarchical clustering heatmaps for the top expression correlates of epigenetic age acceleration.** The dendrograms represent the liver expression of **(a)** mRNA, and **(b)** proteins that are correlated with age acceleration derived from both the pan-tissue general clock, and the liver interventional clock.

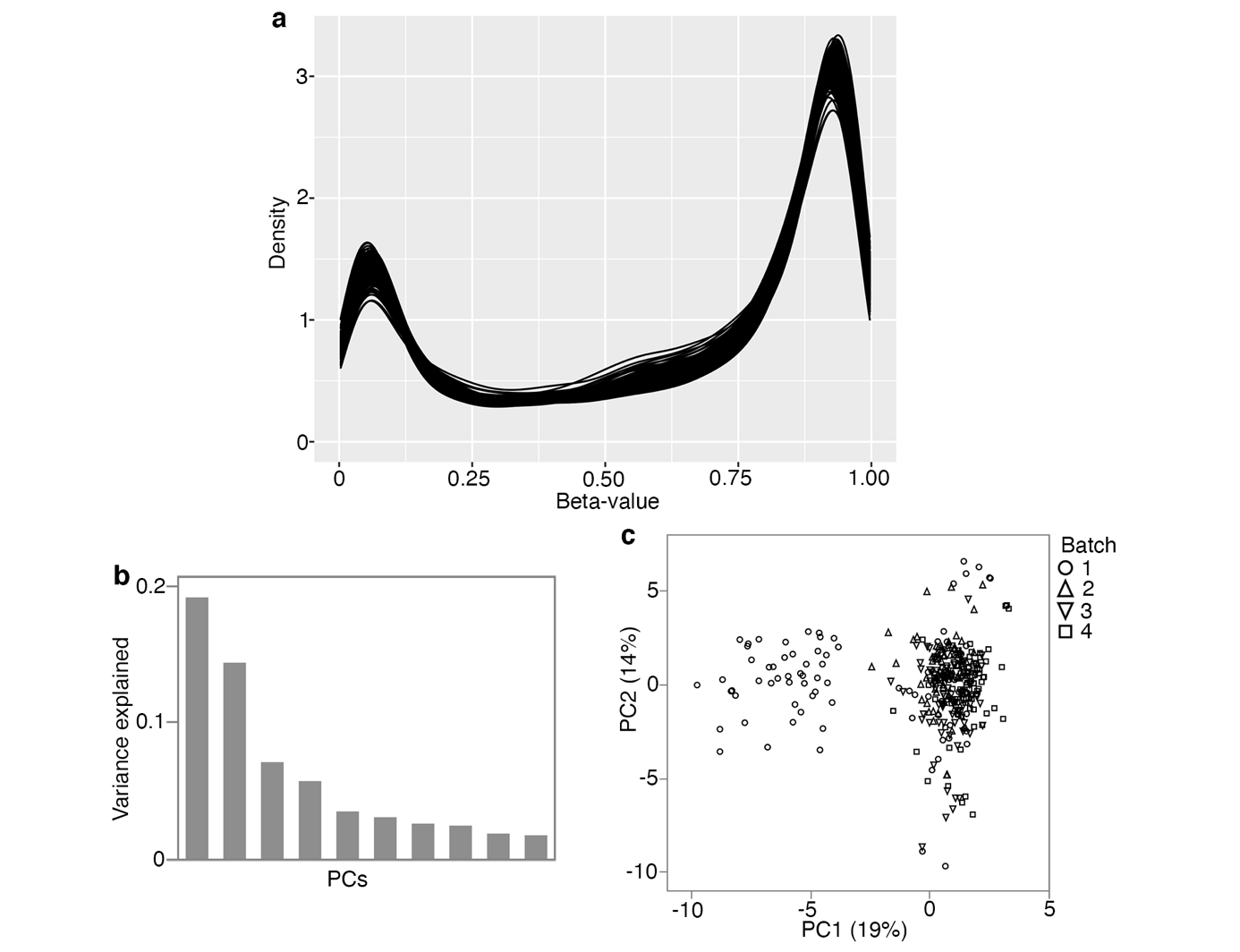

**Fig S6.** **Array quality check**

**(a)** Density plot for the 339 cases using the full set of CpG probes. **(b)** Variance explained by the top 10 principal components (PCs) derived from the full set of probes. **(c)** Plot between component 1 and 2 shows the PC1 captures some batch effect. Here batch is the 96-well plates.

**Supplementary Tables and Figures**

**Table S1. Sex differences in epigenetic aging after correction for body weight**

| **Predictor** | **Outcome** | **Estimate** | **Std Error** | **t Ratio** | **p** |
| --- | --- | --- | --- | --- | --- |
| Sex[M]  lm(x ~ sex + BWF)  n = 58  (40 females, 18 males) | EAA, pan | 0.171 | 0.066 | 2.61 | 0.012 |
|  | EAA, liver | 0.098 | 0.055 | 1.79 | 0.079 |
|  | dev.EAA, pan | 0.180 | 0.059 | 3.05 | 0.004 |
|  | dev.EAA, liver | 0.176 | 0.057 | 3.07 | 0.003 |
|  | int.EAA, pan | 0.324 | 0.077 | 4.20 | 9.9E-05 |
|  | int.EAA, liver | 0.133 | 0.078 | 1.72 | 0.092 |

**Table S2. Pearson correlations between epigenetic age acceleration and strain-level longevity summaries**

| **DNAm** | **minLS** | | **25Q-LS** | | **Mean LS** | | **Median LS** | | **75Q-LS** | | **Max LS** | |
| --- | --- | --- | --- | --- | --- | --- | --- | --- | --- | --- | --- | --- |
|  | **r** | **p** | **r** | **p** | **r** | **p** | **r** | **p** | **r** | **p** | **r** | **p** |
| EAA, pan | 0.08 | 0.15 | 0.06 | 0.27 | -0.06 | 0.34 | -0.08 | 0.17 | -0.16 | 0.005 | -0.13 | 0.02 |
| EAA, liver | -0.02 | 0.73 | -0.03 | 0.58 | -0.07 | 0.24 | -0.08 | 0.18 | -0.08 | 0.15 | -0.06 | 0.27 |
| dev.EAA, pan | -0.02 | 0.70 | 0.03 | 0.59 | -0.02 | 0.69 | -0.04 | 0.50 | -0.06 | 0.33 | -0.01 | 0.91 |
| dev.EAA, liver | -0.05 | 0.35 | -0.01 | 0.83 | -0.05 | 0.37 | -0.08 | 0.18 | -0.07 | 0.21 | 0.01 | 0.86 |
| int.EAA, pan | 0.02 | 0.73 | -0.01 | 0.91 | -0.05 | 0.35 | -0.09 | 0.12 | -0.08 | 0.17 | -0.04 | 0.48 |
| int.EAA, liver | -0.19 | 0.0007 | -0.15 | 0.01 | -0.19 | 0.001 | -0.19 | 0.001 | -0.18 | 0.002 | -0.14 | 0.02 |

n = 302 females BXDs belonging to 65 genotype-by-diet combinations with at least n = 5 observations of age at natural death.

38 BXD genotypes on normal chow, and 27 BXD genotypes on high fat diet.

**Table S3. High priority candidate genes modulating rates of epigenetic aging**

|  |  | |  | **Missense/stop variant in the BXD** | | | |  | **Human GWAS** | |
| --- | --- | --- | --- | --- | --- | --- | --- | --- | --- | --- |
| **Gene** | | **Chr** | **Mb** | **Csq ^a^** | **dbSNP ^a^** | **Ref ^a^** | ***D* ^a^** | **LOD (high allele)^b^** | **Human trait** | **Reference** |
| *Mmd* | | 11 | 90.25 | cis-eQT |  |  |  | 7.4 (B) | Menarche (age at onset) | ^1^ |
| *Stxbp4* | | 11 | 90.54 | missense; cis-eQT | rs3668623 | C | T | 6.1 (D) | Intrinsic epigenetic age acceleration; Menarche (age at onset) | ^2-4^ |
| *Tom1l1* | | 11 | 90.67 | missense; cis-eQT | rs13469307; rs13469308 | G; G | A; C | 2.7 (D) | Parental longevity (mother's age at death or mother's attained age) | ^5^ |
| *Abi3* | | 11 | 95.83 | missense | rs29392269 | G | A |  | Late onset & and family history of Alzheimer's disease | ^6,7^ |
| *Cdk12* | | 11 | 98.20 | missense | rs27086373 | C | A |  | Menopause (age at onset) | ^8^ |
| *Cyp26a1* | | 19 | 37.70 | missense | rs8236989 | G | A |  | Human longevity | ^9^ |
| *Myof* | | 19 | 38.00 | missense | rs31052565; rs46477910 | A; G | G; T |  | Human longevity | ^9^ |
| *Ccnj* | | 19 | 40.83 | missense; cis-eQT | rs36487301 | C | A | 4.1 (D) | Menopause (age at onset) | ^10^ |
| *Nkx2-3* | | 19 | 43.62 | missense | rs30898786 | T | G |  | Epigenetic age acceleration (PhenoAge) | ^11^ |
| *Cutc* | | 19 | 43.75 | cis-eQT |  |  |  | 37.6 (B) | Epigenetic age acceleration (Hannum and PhenoAge) | ^11^ |
| *Chuk* | | 19 | 44.08 | missense | rs48727905 | T | G |  | Age at menopause | ^1^ |
| *Pkd2l1* | | 19 | 44.15 | missense | rs30956598; rs13483639 | A; T | G; C |  | Parental extreme longevity (95 years and older) | ^12^ |

**^a^** Data from the Wellcome Sanger Institute Mouse Genome Project. Each gene also contains multiple non-coding variants. Ref (reference) is the allele for C57BL/6J; *D* is the allele for DBA/2J.

**^b^** LOD score for gene expression *cis*-eQTL in BXD liver. *B* means the C57BL/6J allele has the positive additive effect; *D* means the DBA/2J allele has the positive additive effect.
